# Enhanced mirroring upon mutual gaze: Multimodal evidence from TMS-assessed corticospinal excitability and the EEG mu rhythm

**DOI:** 10.1101/2020.03.24.995696

**Authors:** Jellina Prinsen, Kaat Alaerts

## Abstract

Eye-to-eye contact is a salient cue for regulating everyday social interaction and communication. Previous research has demonstrated that direct eye contact between actor and observer specifically enhances the ‘mirroring’ of others’ actions in the observer, as measured by transcranial magnetic stimulation (TMS)-induced motor evoked potentials (MEPs; an index of motor cortex excitability during action observation). However, it remains unknown whether other markers of mirror system activation, such as suppression of the EEG mu rhythm (i.e. attenuation of neural oscillations in the 8-13 Hz frequency band over the sensorimotor strip), are also susceptible to perceived eye contact. In the current study, a multimodal approach was adopted to assess both TMS-induced MEPs and EEG mu suppression (in separate sessions), while 32 participants (20 men; mean age: 24;8 years) observed a simple hand movement in combination with direct or averted gaze from the live stimulus person. Both indices of mirror system functioning were significantly modulated by perceived eye gaze; showing a significant increase in MEP amplitude and a significant attenuation of the mu rhythm when movement observation was accompanied with direct compared to averted gaze. Importantly, while inter-individual differences in absolute MEP and mu suppression scores were not significantly related, a significant association was identified between gaze-related changes in MEP responses and mu suppression. As such, it appears that while the neurophysiological substrates underlying mu suppression and TMS-induced MEP responses differ, both are similarly affected by the modulatory impact of gaze-related cues. In sum, our results suggest that both EEG mu rhythm and TMS-induced MEPs are sensitive to the social relevance of the observed actions, and that a similar neural substrate may drive gaze-related changes in these distinct markers of mirror system functioning.

## 1 Introduction

Ever since the discovery of ‘mirror neurons’ in the macaque brain, firing not only when the monkey executes a motor action, but also when the monkey merely observes another individual performing that action (di Pellegrino, Fadiga, Fogassi, Gallese, & Rizzolatti, 1992), the description of a homologous action observation-execution matching or ‘mirror system’ and its properties in humans has been a topic of increasing interest. Accordingly, a variety of neuroimaging and electrophysiological techniques – including functional magnetic resonance imaging (fMRI), magneto- or electroencephalography (M/EEG), and transcranial magnetic stimulation (TMS) – have been adopted to identify patterns of mirror system activity during movement observation.

One commonly used method is TMS, a non-invasive brain stimulation technique that activates cortical neurons via the administration of a brief magnetic pulse to the scalp. When TMS is administered over the somatotopically organized primary motor cortex (M1), it induces an involuntary muscle contraction or motor evoked potential (MEP) in the corresponding peripheral muscles (measured with electromyography; EMG), of which the peak-to-peak amplitude reflects variations in M1 excitability. In a seminal study, Fadiga, Fogassi, Pavesi and Rizzolatti (1995) showed that TMS-evoked MEP amplitudes within the stimulated muscles are specifically enhanced during the observation of others’ movements. Subsequent TMS studies have confirmed these observations (for a review, see Fadiga, Craighero, & Olivier, 2005), and provided evidence that the human observation-to-execution matching mechanism is specific to the muscles recruited in the observed actions (Alaerts, Heremans, Swinnen, & Wenderoth, 2009; Alaerts, Swinnen, & Wenderoth, 2009; Fadiga et al., 1995; Strafella & Paus, 2000), with a close temporal coupling (Gangitano, Mottaghy, & Pascual-Leone, 2001).

Another commonly adopted method for investigating mirror system activity relates to the assessment of oscillations in the EEG-based mu rhythm, usually defined in the 8-13 Hz frequency band and topographically centered over the sensorimotor regions of the brain (i.e. electrode positions C3, Cz, and C4 according to the 10-20 international system of electrode placement). At rest, sensorimotor neurons fire in synchrony, leading to high mu power. When a person performs, observes or imagines themselves performing an action, the firing of these neurons has been shown to become increasingly desynchronized, leading to a task-induced suppression of the mu rhythm (Muthukumaraswamy, Johnson, & McNair, 2004; Pfurtscheller, Brunner, Schlögl, & Lopes da Silva, 2006, see Fox et al., 2016 for a meta-analysis). The notion that decreased mu power is related to sensorimotor activation received overall support from EEG-fMRI studies, showing a negative relationship between mu power and the BOLD signal in brain areas considered part of the mirror system (Arnstein, Cui, Keysers, Maurits, & Gazzola, 2011; Braadbaart, Williams, & Waiter, 2013; Perry & Bentin, 2009). Also several MEG studies, having superior spatial resolution compared to EEG, have shown that sensorimotor cortices are significantly modulated by action observation and execution (Hari et al., 1998; Järveläinen, Schürmann, Avikainen, & Hari, 2001).

While the exact role of the mirror system in human social cognition is still a matter of debate, it is generally assumed that the simulation of observed movements in the observer’s own motor system contributes to action recognition and understanding, including related socio-cognitive processes that are important for everyday social interaction such as imitation, mimicry, motor planning and gestural performance (Rizzolatti & Craighero, 2004). The mirror system has also been implicated to be involved in higher-order mentalizing processes, such as inferring others’ intentions (for a review, see Rizzolatti & Fabbri-Destro, 2008; and specific studies by Becchio et al., 2012; Iacoboni et al., 2005), as well as empathy (a form of ‘emotional’ imitation; Iacoboni, 2009). One highly powerful cue for driving interpersonal communications and for conveying (social) intentions is eye contact. Whereas perceived direct gaze from others is indicative of their communicative intent, observed averted gaze signals that the attention of others is directed elsewhere. Accordingly, perceiving the gaze of others has been shown to influence several socio-cognitive processes and behavioral responses in the observer (for relevant reviews, see Conty, George, & Hietanen, 2016; Hietanen, 2018; Senju & Johnson, 2009). In terms of mirror system activity, several TMS studies have shown that under various experimental conditions, perceived communicative intent from the actor (conveyed by different gaze cues) significantly modulates M1 excitability (MEP responses) in the observer (Betti et al., 2019; Prinsen & Alaerts, 2019; Prinsen et al., 2017; Prinsen, Brams, & Alaerts, 2018). To date however it remains unexplored whether suppression of the EEG mu rhythm upon action observation is similarly modulated by social context or communicative intent (i.e. as conveyed by eye contact).

In this respect, it is worth noting that while both TMS and EEG techniques have been widely adopted to investigate observation-to-execution mapping processes, the direct relationship between facilitation of M1 excitability (as assessed with TMS) and suppression of mu rhythm (assessed with EEG) upon movement observation is not well established. In one study, Andrews, Enticott, Hoy, Thomson and Fitzgerald (2015) revealed a significant positive relationship between concurrent recordings of observation-induced mu suppression and M1 excitability in schizophrenia patients and healthy controls. In contrast however, Lepage, Saint-Amour and Théoret (2008) simultaneously recorded M1 excitability and mu suppression during movement observation, imagination and execution of simple hand actions in healthy adult participants and showed that while both measures were significantly modulated by the experimental conditions (increased M1 excitability, increased mu suppression), changes in M1 excitability were not significantly correlated to changes in mu suppression at the inter-individual subject level. Similarly, in two recent studies assessing observation-induced changes in M1 excitability and mu suppression, either simultaneously (Cole, Barraclough, & Enticott, 2018) or in different recording sessions (Lapenta, Ferrari, Boggio, Fadiga, & D’Ausilio, 2018), no direct relationship was revealed between the two measures. Accordingly, it has been suggested that M1 excitability and mu suppression may represent different aspects of the mirror system, presumably due to the different spatial and temporal characteristics of the two techniques. Indeed, while EEG mu suppression indexes the sum of post-synaptic neuronal activity over a large area (not restricted to M1) over a relatively long time period (typically > 1 second), TMS assesses changes in M1 excitability by stimulating a relatively small population of neurons (at the level of M1) at a discrete point in time (Andrews et al., 2015; Pineda, 2005; Rossini et al., 1994).

Within the present study we adopted a multi-modal approach for assessing gaze-related modulations of mirror system activity, by recording - from the same participant sample - indices of TMS-induced MEP responses and EEG mu rhythm suppression upon movement observation with variable communicative intent. In particular, the two mirror system indices were recorded in separate sessions while participants observed simple intransitive hand movements accompanied by either direct gaze (signaling communicative intent) or averted gaze from the stimulus person. Based on the findings of Prinsen and Alaerts (2019), and following the recommendations by Reader and Holmes (2016), a naturalistic two-person paradigm was adopted, incorporating a ‘live’ stimulus person to convey the gaze and movement cues. In line with previous TMS studies demonstrating an effect of observed eye contact on the mirroring of others’ actions (Prinsen & Alaerts, 2019; Prinsen et al., 2017, 2018), it was hypothesized that TMS-induced MEPs are enhanced upon movement observation accompanied with direct eye gaze, compared to movement observation accompanied with averted gaze. A key question was to assess whether suppression of the mu rhythm is also susceptible to a top-down response modulation by perceived communicative intent (i.e. eye contact); and whether eye contact-induced changes in M1 excitability are associated with changes in EEG mu suppression. Lastly, since the central mu rhythm oscillates in the same 8-13 Hz frequency band and displays similar response properties as occipital alpha rhythms (i.e. dominant when at rest, suppressed by perceptual events and attentional processing), an important issue in EEG action observation studies is the potential contamination of the mu rhythm by changes in alpha. In line with recent guidelines by Fox et al. (2016) and Hobson and Bishop (2017), occipital alpha suppression was also taken into account.

## 2 Method and Materials

### 2.1. Participants

A total of 32 individuals (20 men and 12 women) aged between 18 and 36 years old (mean ± SD: 22;9 ± 3;7 years; months) participated in this study. All participants were right-handed, which was confirmed with the Edinburgh Handedness Questionnaire (EHQ; Oldfield, 1971). Exclusion criteria comprised medication use, any diagnosed psychiatric (e.g. ASD, ADHD) or neurological disorder (e.g. stroke, epilepsy, concussion), left handedness or any contraindication for TMS (Rossi, Hallett, Rossini, & Pascual-Leone, 2012). Written informed consent was obtained from all participants prior to the experimental procedure. Ethical approval for the experimental protocol was granted by the local Ethics Committee for Biomedical Research at the University of Leuven in accordance to the Declaration of Helsinki (World Medical Association, 2013).

### 2.2. Experimental protocol and stimuli

Participants were seated at a distance of approximately 80 cm from a 20 × 30 cm voltage-sensitive liquid crystal (LC) shutter screen (DreamGlass Group, Spain) attached to a black frame and were instructed to observe and pay close attention to the presented stimuli. A ‘live’ female stimulus person (experimenter J.P.) was seated behind the panel (similar set-up as Prinsen & Alaerts, 2019). During the experimental conditions, the stimulus person’s face was presented through the LC shutter screen for 4 seconds. Importantly, the stimulus person was unknown to the participants and only briefly interacted with them before the experimental procedure. While the LC screen was transparent, the stimulus person either gazed directly towards the observing participant (i.e. engaging in mutual eye contact) or displayed a gaze 30° to the right (i.e. showing averted gaze). During both gaze conditions, the stimulus person held her right hand horizontally beneath her face with the dorsal side directed to the participants and performed a simple index finger abduction movement. The stimulus person bore a neutral expression and tried to avoid eye blinks during the duration of the trial. An illustration of the experimental conditions is provided in Prinsen and Alaerts (2019).

Each gaze condition was presented 20 times in 4 second trials with an inter-stimulus-interval of 2 seconds, during which the shutter remained opaque (similar set-up as Prinsen & Alaerts, 2019). The same stimulus protocol was adopted for the TMS and EEG assessment. In order to ensure that all participants viewed and attended the stimuli properly, participants were asked once at a random point in time during each neurophysiological assessment (in-between trials) to verbally report the stimulus that was presented in the previous trial. Participants were able to correctly report the presented stimulus in 98.5% of the assessments, indicating that they attended the stimuli properly.

Mirror system activity was investigated in two assessment sessions conducted on the same day, with a fifteen minute break between sessions. In one session, stimulus presentation was accompanied with transcranial magnetic stimulation (TMS) in order to assess excitability at the level of the primary motor cortex. In the other session, electroencephalography (EEG) assessments were performed in order to measure mu rhythm suppression. The order of assessment method (TMS or EEG) was counterbalanced across participants.

### 2.3. Neurophysiological assessment

#### 2.3.1. TMS and EMG recordings

The TMS and EMG electrode set-up is illustrated in **figure 1A**. During observation of the stimuli, single-pulse TMS was administered over the primary motor cortex (M1) with a hand-held 70 mm figure-of-eight coil (oriented approximately 45° relative to the mid-sagittal line) and a Magstim-200 stimulator (Magstim Company Ltd., UK). Optimal coil location for the experimental TMS-stimulation was determined as the site that produced maximal responses (i.e. MEPs) while at rest (“hotspotting”) in the contralateral first dorsal interosseous (FDI) muscle, a muscle implicated in the to-be-observed index finger opening movement. Resting motor thresholds (rMT) were individually defined as the lowest stimulation intensity that produced a peak-to-peak MEP of at least 50 µV in five out of ten consecutive trials (Rossini et al., 1994). Experimental stimulation intensity was set at a supra-threshold of 130% of the subject’s rMT. In each trial, a single TMS pulse was delivered on the third second of stimulus presentation, which coincided with the execution of the index finger opening movement of the stimulus person.

**Figure 1.**
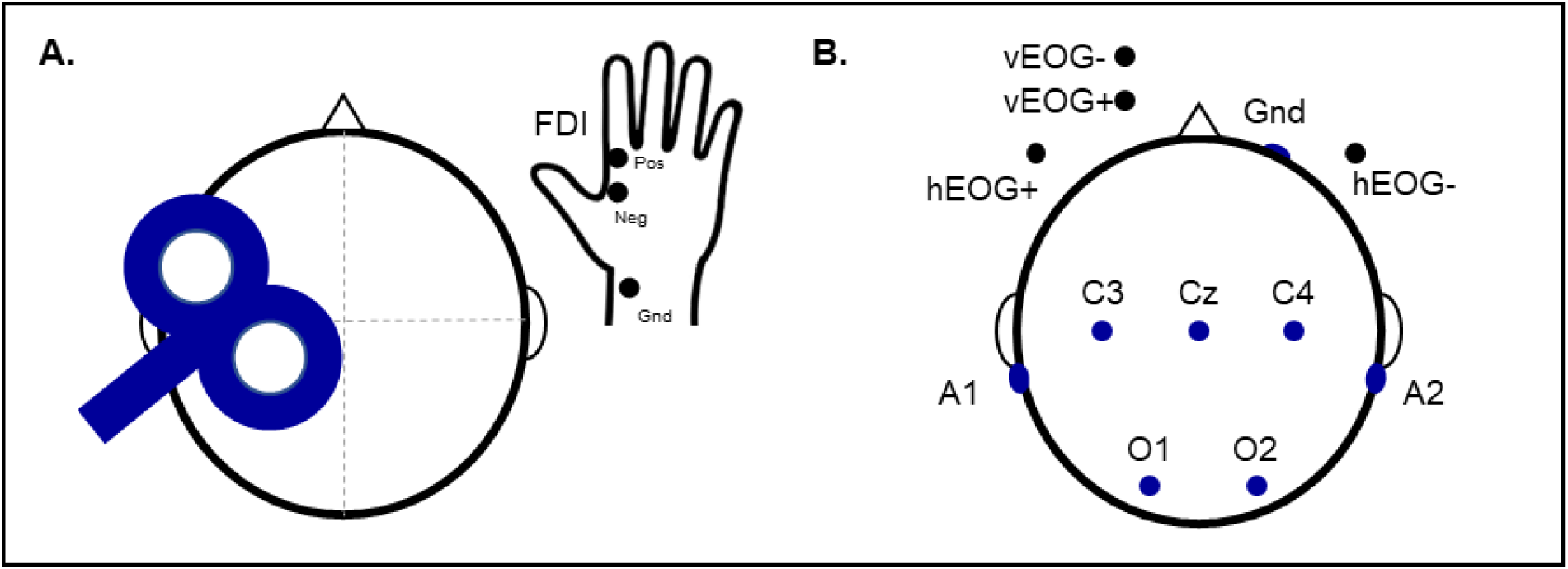
**(A)** MEPs induced by TMS over the left primary motor cortex were recorded from EMG electrodes located on the FDI index finger muscle of the right hand. **(B)** Continuous EEG was acquired from electrode sites C3, Cz and C4 to calculate mu suppression, and sites O1 and O2 for alpha suppression.

Surface electromyography (EMG) recordings were obtained using disposable adhesive electrodes arranged in a tendon-belly montage. The EMG signal was sampled at 2 kHz, band-pass filtered (5-1000 Hz) and analyzed offline. Signal software (version 6.02, Cambridge Electronic Design, UK) and a CED Power 1401 analog-to-digital converting unit (Cambridge Electronic Design, UK) were used for EMG-recordings, triggering of the TMS-stimulator and shifting of the LC window from an opaque to transparent state.

#### 2.3.2. EEG data acquisition

The NeXus-32 multimodal acquisition system and BioTrace+ software (version 2015a, Mind Media, The Netherlands) were used to collect electroencephalography (EEG) recordings. Continuous EEG was recorded with a cap with 22 sintered Ag/AgCL embedded electrodes (MediFactory, The Netherlands), incorporating 19 EEG channels configured according to the international 10-20 system of electrode placement, two reference electrodes located on the left and right mastoid bones behind the ear (A1 and A2), and a AFz ground electrode. The EEG signal was amplified using a unipolar amplifier and mathematically referenced offline to linked mastoids. Gentle skin abrasion and electrode paste (combination of electrolytic NuPrep gel and conductive 10-20 paste) were used to reduce electrode impedances below 10 kΩ. Eye movements as well as eye blinks were monitored using two pairs of bipolar electro-oculogram (EOG) electrodes, one pair attached to the external canthi of each eye (horizontal eye movements; hEOG) and one pair attached below and above the left eye (vertical eye movements; vEOG). The sampling rate of the recordings was 256 Hz. E-Prime 2.0 software (Psychology Software Tools Inc., USA) and the NeXus Trigger Interface (NTI, 2048 Hz sample rate; Mind Media, The Netherlands) were used to synchronize stimulus events with the NeXus-32 EEG recordings and the triggering of the LC window.

### 2.4. Data handling and preprocessing

#### 2.4.1. TMS-induced MEPs

Based on the recorded EMG data, peak-to-peak amplitudes of the TMS-induced MEPs were determined using in-house MATLAB scripts (version R2015a, MathWorks Inc., USA). Additionally, background EMG was quantified by calculating the root mean square (RMS) across the 110 to 10 millisecond interval prior to TMS-stimulation. For a given subject, trials with excessive pre-TMS tonic muscle activity (background EMG exceeding 2.5 standard deviations from the mean) were excluded from analysis (1.95% of trials). Trials with extreme MEP-amplitudes (exceeding 1.5 interquartile distances from the mean) were also discarded. This resulted in an additional omission of 8.98% of trials. The total number of discarded trials was similar across gaze conditions (all *p* > .11). MEP peak-to-peak amplitudes were log-transformed to conform to normality.

#### 2.4.2. EEG mu/alpha suppression calculation

Two participants were excluded from the final analysis due to technical malfunctions of the NeXus Trigger Interface, used for time-locking EEG data with the stimulus presentation. EEG data of the remaining participants (*n* = 30) were preprocessed and analyzed offline using BrainVision Analyzer 2 software (version 2.2, Brain Products GmbH, Germany). The raw EEG signal was filtered using a 0.5 - 40 Hz IIR band-pass filter with zero phase shift (Butterworth, 24 dB). Taking into account the vEOG and hEOG channels, deflections resulting from eye blinks and horizontal eye movements were removed by the implemented Independent Component Analysis (ICA) module in BrainVision Analyzer 2 (Jung et al., 2000). Cleaned EEG data were segmented separately for each condition into segments of 4 seconds, corresponding to the duration of the trial. Segments with residual artifacts exceeding ± 100 µV in amplitude were rejected. Note that one additional participant was removed from the final analyses due to excessive artifacts (> 60% of artifactual segments). For each segment, the spectral power (µV^2^) in the 8-13 Hz range was computed using the Fast Fourier Transform (FFT; including a Hanning window with an attenuation domain of 25%). Obtained power values were then averaged separately for each experimental gaze condition and electrode.

Suppression indices were computed at three central sites (C3, Cz and C4) located over the sensorimotor strip where mu rhythm modulations are expected. To assess the spatial specificity of the gaze-dependent modulations in the mu rhythm, alpha suppression indices were also calculated for occipital electrodes O1 and O2 (**figure 1B**). Mu and alpha suppression indices for each electrode were calculated as the log-transformed ratio of the 8-13 Hz band power during the 4-second trials relative to the power of a 1-second interval prior to the start of the trial (baseline). Log ratios lower than zero indicate suppression.

### 2.5. Data analysis and statistics

In order to investigate eye contact-induced changes in M1 excitability, a repeated measures analysis of variance (RM-ANOVA) with within-subject factor ‘observed gaze’ (averted, direct) was performed on the MEP peak-to-peak amplitude data. For the EEG data; it was first tested whether all movement observation conditions elicited a significant suppression relative to the pre-trial baseline segments (as recommended by Hobson & Bishop, 2017); i.e. ratio values were tested using single-sample *t*-tests against a value of 0, separately for each gaze condition (**table 1**). The mu and alpha suppression indices were analyzed separately using a RM-ANOVA with the within-subject factors ‘observed gaze’ (averted, direct) and ‘electrode’ (mu: C3, Cz and C4; alpha: O1 and O2). For the RM-ANOVAs, the categorical factor ‘session’ was included as effect-of-no-interest to account for potential effects of counter-balancing the two assessment sessions (i.e. TMS or EEG first). Significant interaction effects were further investigated by means of Fisher Least Significant Differences (LSD) post-hoc tests, the partial Eta square (*η*^*2*^_*p*_) value was calculated as an estimate of effect size.

**Table 1.** Mean central mu and occipital alpha suppression (in 8-13 Hz frequency band) for each gaze condition (averted, direct) and electrode. \*\**p* < .001

|  | Averted gaze |  | Direct gaze |  |
| --- | --- | --- | --- | --- |
| | Mean (SD) | $t$ against 0 | Mean (SD) | $t$ against 0 |
| **Central electrodes** |  |  |  |  |
| C3 | -0.661 (0.131) | -26.71** | -0.684 (0.106) | -33.98** |
| Cz | -0.711 (0.128) | -29.78** | -0.758 (0.134) | -30.33** |
| C4 | -0.687 (0.121) | -30.69** | -0.711 (0.091) | -42.02** |
| **Occipital electrodes** |  |  |  |  |
| O1 | -0.762 (0.176) | -22.91** | -0.781 (0.177) | -23.32** |
| O2 | -0.744 (0.177) | -22.32** | -0.748 (0.186) | -21.19** |

In order to directly investigate the relationship between the effects of eye contact on the different measures, the ‘eye contact effect’ was quantified for each subject as the percentage change (%change) in the direct gaze condition relative to the averted gaze condition (similar approach as Enticott, Kennedy, Bradshaw, Rinehart, & Fitzgerald, 2011): 

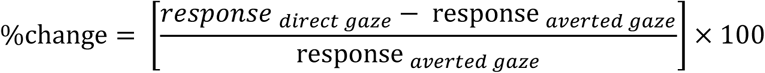

Higher mirror responses during the direct versus averted gaze condition are indicated by a positive %change score for MEPs, and a negative %change score for mu and alpha suppression indices. Pearson correlation analyses were performed to assess the association between the measures. For all performed correlations, the Cook’s distance metric was used to identify influential data points (defined as Cook’s *D* > 1), but none were detected. The coefficient of determination (*R*^*2*^) is reported as an estimate of effect size. All statistics were calculated with Statistica 10 (StatSoft, USA). Results were considered significant when *p* < .05.

## 3 Results

### 3.1. MEP results

A RM-ANOVA with within-subject factor ‘observed gaze’ (averted, direct) was performed on the MEP data to investigate the effect of observed gaze on M1 excitability. The mean (log-transformed) MEP peak-to-peak amplitudes for each gaze condition is presented in **figure 2A**. In line with our hypothesis, a significant main effect of perceived gaze direction (*F*(1,31) = 9.53, *p* = .005, *η*^*2*^_*p*_ = .24) was revealed. Thus, in accordance with previous TMS studies investigating the eye contact effect on M1 excitability (Prinsen & Alaerts, 2019; Prinsen et al., 2017, 2018), MEPs recorded from the FDI muscle were significantly higher when movement observation was accompanied with direct eye gaze from the stimulus person (mean: 0.231, SD: 0.356), compared to averted gaze (mean: 0.161, SD: 0.389). Note that no significant main or interaction effects were found for the categorical ‘session’ factor-of-no-interest (all *p* > .42).

**Figure 2.**
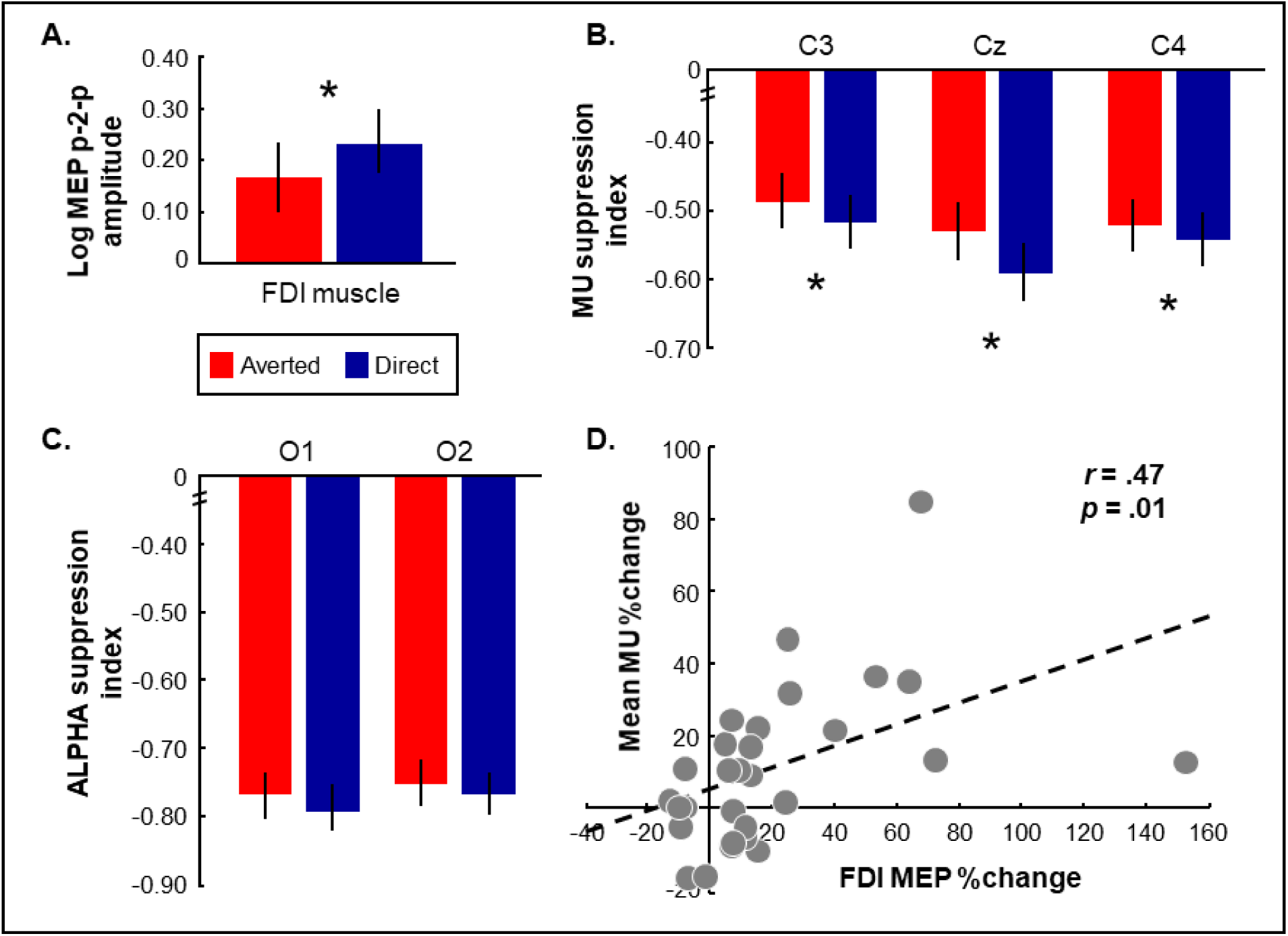
**(A)** Significant effect of perceived eye gaze (direct, averted) on log-transformed MEP peak-to-peak amplitude scores recorded from the FDI muscle **(B)** and mu suppression indices per central electrode. \**p <* .05, vertical error bars denote mean ± SE. **(C)** The effect of perceived eye gaze on alpha suppression over occipital electrodes was not significant. **(D)** A significant positive correlation was found between condition-specific (i.e. eye contact related) changes in MEP amplitude and EEG mu suppression (averaged across central electrodes).

### 3.2. EEG results

Significant decreases in mu power with respect to the included rest condition were encountered for each electrode and gaze condition (single sample *t*-tests against 0: all *p* < .001; see **table 1**), signaling that all gaze conditions induced an overall significant suppression of the mu rhythm during movement observation in the central electrodes.

A RM-ANOVA with observed gaze condition (averted, direct) and electrode (C3, Cz, C4) as within-subject factors revealed a significant main effect of observed eye gaze (*F*(1,26) = 6.97, *p* = .01, *η*^*2*^ = .21), but no gaze × electrode interaction (*F*(2,52) = 2.72, *p* = .07, *η*^*2*^_*p*_ = .09). This shows that, irrespective of electrode, mu rhythm suppression upon movement observation was more pronounced during the direct versus the averted gaze condition (see **figure 2B**). Also a significant main effect of electrode was revealed (*F*(2,52) = 6.81, *p* = .002, *η*^*2*^_*p*_ = .21), indicating that, irrespective of gaze condition, mu rhythm suppression was slightly more pronounced in electrode Cz (see **figure 2B** and **table 1**). Note that no significant main or interaction effects were found for the categorical ‘session’ factor-of-no-interest (all *p* > .08).

Also alpha activity from occipital electrodes O1 and O2 was significantly suppressed during all gaze conditions compared to rest (all *p* < .001; see **table 1**). Importantly however, a comparable RM-ANOVA as described above did not indicate a significant main effect of observed eye gaze (*F*(1,25) = 1.90, *p* = .18, *η*^*2*^_*p*_ = .07) or electrode (*F*(1,25) = 1.53, *p* = .23, *η*^*2*^_*p*_ = .06) for these occipital electrodes, nor an electrode × gaze interaction (*F*(1,25) = 0.08, *p* = .78,, *η*^*2*^_*p*_ = .003) (**figure 2C**). During movement observation, occipital alpha suppression was thus not significantly modulated by the observed gaze cues.

### 3.3. TMS-EEG correlations

Exploration of a potential relationship between FDI MEP amplitudes and mu suppression scores over the central electrodes (averaged score) revealed no significant correlation between absolute MEP and mu responses for either the direct gaze condition (*r*(29) = -.18, *p* = .36, *R*^2^ = .03), nor for the averted gaze condition (*r*(29) = -.15, *p* = .43, *R*^*2*^ = .03). Note that the absence of a significant association persisted when only mu suppression over electrode C3 (contralateral to right-hand MEPs and corresponding to the site of TMS stimulation) was considered (averted gaze: *r*(28) = -.06, *p* = .73, *R*^2^ = .005; direct gaze: *r*(28) = .01, *p* = .95, *R*^2^ < .01)

Interestingly however, it was shown that for the experimental ‘eye contact effect’ (see section 2.5 in Method and Materials), 22% of the variance was shared between the TMS and EEG measures, indicating that increments in MEP amplitude in response to direct gaze were significantly associated with similar enhancements of mu suppression (*r*(29) = .47, *p* = .01, *R*^2^ = .22; **figure 2D**). Importantly, this association was specific to the central electrodes, as MEPs were not significantly correlated to alpha suppression indices over occipital electrodes, either in terms of absolute responses (averted gaze: *r*(29) = -.21, *p* = .26, *R*^2^ = .04; direct gaze: *r*(29) = -.19, *p* = .32, *R*^*2*^ = .04), or in terms of the experimental ‘eye contact effect’ (*r*(29) = .15, *p* = .43, *R*^*2*^ = .02).

## 4 Discussion

This study aimed to investigate the impact of observed gaze cues on TMS- and EEG-based measures of mirror system activity. In agreement with previous studies (Prinsen & Alaerts, 2019; Prinsen et al., 2017, 2018), we showed that motor cortex (M1) excitability assessed as TMS-induced MEPs upon movement observation was significantly impacted by observed gaze direction from the observed stimulus person. Furthermore, we demonstrated – for the first time – that also EEG-based mu suppression (in the 8-13 Hz frequency band) over the sensorimotor strip (electrodes C3, Cz, C4) was enhanced when observing direct, compared to averted eye gaze from the actor. Importantly, while absolute MEP and mu suppression scores were not significantly related, a significant association was identified between gaze-related changes in M1 excitability and mu suppression.

The observation that both TMS-induced MEPs and EEG-based mu suppression are modulated by observed gaze cues is in line with the recent notion that motor resonance is not a static process, but is adapted depending on the social context in which the observed movements are embedded. According to these theoretical proposals, this subtle social top-down control of motor resonance processes is proposed to originate in the mentalizing system (Vogeley, 2017; Wang & Hamilton, 2012; Yang et al., 2015). Other TMS studies have indicated that M1 excitability is also modulated by other social factors, such as, amongst others, emotional body language of the actor (Borgomaneri, Vitale, Gazzola, & Avenanti, 2015), social reciprocity (Sartori, Cavallo, Bucchioni, & Castiello, 2012), and the level of observed social interaction (Donne, Enticott, Rinehart, & Fitzgerald, 2011; Hogeveen & Obhi, 2012). Similarly, EEG activity in the mu frequency range has been demonstrated to depend on the extent by which participants are engaged in a social game (Perry, Stein, & Bentin, 2011), the perception of social information such as intentions and emotions (Perry, Troje, & Bentin, 2010) and empathic processes (Hoenen, Schain, & Pause, 2013). Taken together, this subtle and sophisticated adjustment of motor resonance according to the demands of the social context forms an essential competence of humans for flexibly engaging in interpersonal social interactions (Wang & Hamilton, 2012).

A second objective of the current research was to further disentangle the relationship between TMS- and EEG-based measures of mirror system functioning at the inter-individual subject level, as previous studies (Andrews et al., 2015; Cole et al., 2018; Lapenta et al., 2018; Lepage et al., 2008) provided an unclear pattern of results. In terms of absolute responses, we were unable to establish an association between these two measures. This is in accordance with several previous studies who have directly compared mu suppression and M1 excitability in healthy participants, either by adopting passive observation of simple hand actions (Lepage et al., 2008) or goal-directed grasping movements (Lapenta et al., 2018). One additional study, incorporating a mentalizing task to infer others’ intentions in adults with and without an autism spectrum disorder also failed to demonstrate a relationship between these measures (Cole et al., 2018).

On the one hand, this lack of a significant association between absolute mu suppression scores and MEP responses may relate to the substantial differences in neurophysiological underpinnings and temporo-spatial properties between these measures (i.e. induced activation of a small population of M1 neurons recorded at peripheral muscles at a discrete time point versus summed post-synaptic electrical activity from a broad population of sensorimotor neurons over a relatively long time period). In other words; MEP recordings of mirror system functioning during TMS are obtained at the level of the muscle, reflecting corticospinal processes, whereas the EEG mu rhythm mainly reflects central cortical activity. Although both techniques have been shown to reliably capture mirror system activation (see reviews by Fadiga, Carighero & Olivier, 2005; Hobson & Bishop, 2017), it has been suggested that - considering these substantial differences in neurophysiological underpinnings - both techniques might target different aspects of the mirror system. In this respect, the neural processes triggered by action observation have been proposed to be layered in several hierarchically organized functional levels (Grafton & Hamilton, 2007; Kilner, Friston, & Frith, 2007). These proposed levels are (i) the muscular level (decoding the pattern of muscle activity necessary to perform the action); (ii) the kinematic level (mapping the effector movement in time and space); (iii) the aim level (including transitive or intransitive short-term goals); and (iv) the intention level (regarding the long-term purpose of the action). Without explicitly framing their design or results in this theoretical structure, Cole et al. (2018) demonstrated that higher mu suppression was associated with superior mentalizing performances, whereas TMS-induced MEPs showed no differences associated with mentalizing. These findings might suggest that the EEG mu rhythm is able to capture higher-order processes such as intentions, but MEPs are not. Note however that Cole et al. (2018) opted to deliver the TMS pulse *after* the completion of the video clips conveying the intentions of the actor (i.e. not taking the strict temporal coupling for M1 excitability into account). Furthermore, their results contrast a previous study by the same group that investigated mirror system activation during observation of intransitive, transitive and interacting hand movement stimuli in adults with and without schizophrenia (Andrews et al., 2015). This work revealed a positive association between absolute mu suppression and M1 exitability, when averaged across all conditions with biological movement. In sum, future work is necessary to obtain complementary information with respect to this hierarchical organization in terms of absolute responses.

Interestingly, while no direct associations were evident between absolute mu suppression scores and MEPs, it was shown that direct gaze-induced increments in MEP amplitude were paralleled by similar enhancements of mu suppression, as indicated by a significant positive relationship of moderate strength between the ‘eye contact effect’ in the EEG and TMS measures. The relationship between gaze-related changes in both measures is an important finding, since it provides initial indications that the two methods do capture similar flexible changes of these underlying neural processes in response to condition-specific manipulations or contexts (e.g. such as the presentation of socio-communicative cues). These flexible changes across neurophysiological markers can be considered to reflect a similar “gating” mechanism according to the social saliency or relevance of the observed stimuli, whereby the processing of irrelevant stimuli is inhibited in order to better process relevant stimuli (Anderson & Ding, 2011; Kilner, Marchant & Frith, 2006). As such, while the neural correlates underlying absolute MEP and mu suppression scores may not be the same, it appears that the neural regions involved in processing gaze related cues, i.e. superior temporal sulcus (Pelphrey, Viola, & McCarthy, 2004) or associated regions of the mentalizing network (Kampe, Frith, & Frith, 2003), exert a similar modulating impact on the (distinct) neurophysiological substrates that drive mu suppression or TMS-induced MEP responses upon movement observation.

There are several methodological considerations to be taken into account when evaluating the EEG mu rhythm. For an in-depth discussion, the interested reader is referred to reviews by Hobson & Bishop (2017) and Cuevas, Cannon, Yoo, & Fox (2014). Here, we briefly touch upon some relevant issues relevant that motivated our adopted design.

First, given the fact that the mu and alpha rhythms oscillate in the same frequency band and show similar response properties (Hobson & Bishop, 2016, 2017)(Hobson & Bishop, 2016, 2017)(Hobson & Bishop, 2016, 2017)(Hobson & Bishop, 2016, 2017), we also inspected alpha suppression at the occipital electrodes (O1 and O2). Significant alpha suppression was present during movement observation, suggesting that an attentional component might have been at play during the observation of the different stimuli (see also Perry et al., 2011). It is however important to note that, in contrast to the central mu rhythm, the occipital alpha rhythm was not subjected to gaze-related modulations (i.e. alpha suppression was not significantly stronger during direct vs. averted gaze at occipital electrodes). Furthermore, only eye-contact induced changes in mu suppression indices, but not alpha suppression indices, were significantly associated with eye-contact induced changes in MEPs. In this respect we believe that the aforementioned observations highlight the specificity of the mu rhythm in reflecting action-specific mirroring processes, as opposed to reflecting contamination or volume conduction from attentional processes at occipital sites. In line with this notion, Debnath, Salo, Buzzell, Yoo and Fox (2019) recently showed that while both central mu and occipital alpha rhythms are indeed similarly suppressed during movement observation, phase synchrony was only evident between central-occipital areas, but not between neighboring occipital-parietal and central-parietal electrodes. These results exclude the possibility of a general spread of occipital alpha activity due to volume conduction, but suggest that visuospatial attention (indexed by occipital alpha) and sensorimotor mirroring (indexed by central mu) are functionally distinct but highly coordinated processes during action observation (see Debnath et al., 2019 and Fox et al., 2016 for more detailed hypotheses).

Secondly, similar to Lapenta et al. (2018), the current study assessed TMS and EEG related mirror system activity within two separate sessions, whereas the majority of previous studies have used TMS and EEG simultaneously (Andrews et al., 2015; Cole et al., 2018; Lepage et al., 2008). While concurrent recording may allow for a more direct comparison between both indices, the application of magnetic pulses during TMS induces artifacts in the simultaneously recorded EEG signals. It is therefore necessary to specifically exclude the time window that overlaps with the deliverance of the TMS pulse, which is preferably optimized for the action observation scene. As such, some crucial time windows for eliciting mu suppression may be removed.

Lastly, as the key design feature of mu suppression studies is the comparison of an experimental condition to a baseline condition in which one would not expect mirror system activity, the choice of baseline condition has a substantial impact. Ideally, one collects a baseline period just prior to the period of interest (the onset of movement), that is identical to the experimental condition, except for this event of interest (Hobson & Bishop, 2016; Tangwiriyasakul, Verhagen, Van Putten, & Rutten, 2013). However, the associative property of the mirror system might pose difficulties for establishing an optimal baseline condition (note that this is not limited to EEG, but also applies to other modalities in action observation research). Although theoretically speaking mirror system activity would be greatest during movement observation, the mere presence of an interactive agent (or object) may elicit early anticipatory reactivity, especially in a design with multiple repetitions (Cuevas et al., 2014). Indeed, Southgate, Johnson, Osborne and Csibra (2009) have demonstrated anticipatory mu suppression prior to action observation. As few studies to date have taken advantage of the superior temporal resolution of EEG to examine the temporal dynamics of mu suppression, it is important to take into account that changes in mu might take place before, during or after observation of an action (Fox et al., 2016).

## 5 Conclusion

To conclude, both TMS-induced MEPs and EEG-based mu rhythm suppression upon movement observation have independently shown that mirror system activity is significantly impacted by eye contact between observer and performer. Furthermore, this is the first study to date to show that condition-induced (i.e. eye contact-related) changes in M1 excitability and mu suppression are related, providing first evidence that a similar gating mechanism may drive these distinct markers of mirror system functioning.

## 6 Conflict of interest

The authors declare that the research was conducted in the absence of any commercial or financial relationships that could be construed as a potential conflict of interest.

## 7 Acknowledgments

We are thankful for all participating subjects. Furthermore, we would like to thank Sylvie Bernaerts, Nicky Daniels, Elisa Maes, Annelore Deschepper and Julio Rodriguez Larios for their help in conducting the experiment; and Paul Meugens and prof. Stephan P. Swinnen for their methodological and technical support.

## 8 Funding

This research was supported by grants from the Flanders Fund for Scientific Research (FWO [KAN 1506716N, G079017N]) and the Branco Weiss fellowship of the Society in Science - ETH Zurich granted to KA. JP is supported by an internal fund of the KU Leuven [STG/14/001] and the Marguerite-Marie Delacroix foundation. We would also like to thank the Academische Stichting Leuven (2016/131).

